# Modeling the Effects of Transcranial Magnetic Stimulation on Spatial Attention

**DOI:** 10.1101/2023.01.11.523548

**Authors:** Ying Jing, Ole Numssen, Konstantin Weise, Benjamin Kalloch, Lena Buchberger, Jens Haueisen, Gesa Hartwigsen, Thomas R. Knösche

## Abstract

**Objectives:** Transcranial magnetic stimulation (TMS) has been widely used to modulate brain activity in healthy and diseased brains, but the underlying mechanisms are not fully understood. Previous research leveraged biophysical modeling of the induced electric field (E-field) to map causal structure-function relationships in the primary motor cortex. This study aims at transferring this localization approach to spatial attention, which helps to understand the TMS effects on cognitive functions, and may ultimately optimize stimulation schemes.

**Approach:** Thirty right-handed healthy participants underwent a functional magnetic imaging (fMRI) experiment, and seventeen of them participated in a TMS experiment. The individual fMRI activation peak within the right inferior parietal lobule (rIPL) during a Posner-like attention task defined the center target for TMS. Thereafter, participants underwent 500 Posner task trials. During each trial, a 5-pulse burst of 10 Hz repetitive TMS (rTMS) was given over the rIPL to modulate attentional processing. The TMS-induced E-fields for every cortical target were correlated with the behavioral modulation to identify relevant cortical regions for attentional orientation and reorientation.

**Main results:** We did not observe a robust correlation between E-field strength and behavioral outcomes, highlighting the challenges of transferring the localization method to cognitive functions with high neural response variability and complex network interactions. Nevertheless, TMS selectively inhibited attentional reorienting in five out of seventeen subjects, resulting in task-specific behavioral impairments. The BOLD-measured neuronal activity and TMS-evoked neuronal effects showed different patterns, which emphasizes the principal distinction between the neural activity being correlated with (or maybe even caused by) particular paradigms, and the activity of neural populations exerting a causal influence on the behavioral outcome.

**Significance:** This study is the first to explore the mechanisms of TMS-induced attentional modulation through electrical field modeling. Our findings highlight the complexity of cognitive functions and provide a basis for optimizing attentional stimulation protocols.

## 1. Introduction

Transcranial magnetic stimulation (TMS) is a non-invasive brain stimulation technique that allows the modulation of cortical function in vivo. It has been widely used to map structure-function relationships in healthy brains (Bestmann and Feredoes, 2013; Groppa et al., 2013), as well as for therapeutic application (Perera et al., 2016; Rawji et al., 2020). In brief, TMS induces an electric field (E-field) in the brain, which can temporarily excite or inhibit the stimulated area by depolarizing or hyperpolarizing cell membranes (Hallett, 2000). However, the precise location of the neuronal populations which are affected by the induced E-field and cause the observed behavioral or physiological changes are difficult to determine. As a result, TMS studies often exhibit considerable interindividual variability in the observed outcome, which hampers its general efficacy in both basic research and clinical applications (Hartwigsen and Silvanto, 2022). This observed variability in TMS effects may be attributed to a complex interplay of interindividual differences (e.g., tissue conductivity, gyral shape, E-field direction, and magnitude (Numssen et al., 2023) and the variable response of neuronal networks (Hartwigsen and Silvanto, 2022).

To address individual variability in TMS effects, researchers have turned to biophysical modeling of the induced E-field based on individual head anatomy. This approach has been increasingly used to estimate cortical locations and stimulation strengths at target areas (Van Hoornweder et al., 2022; Nieminen et al., 2015; Thielscher et al., 2015). Common methods that consider the biophysical properties of the head models, along with stimulation parameters like intensity, location, and coil orientation, encompass the boundary element method (BEM; Makarov et al., 2021; Weise et al., 2023) and the finite element method (FEM; Thielscher et al., 2015). These approaches provide realistic estimates of the induced E-field distribution in the head. This allows researchers to explore the effects of different stimulation parameters and optimize TMS protocols for specific brain regions and functions. By correlating the E-field with behavioral or physiological outcome measures, one may identify the neural structures that are effectively stimulated and underlie these effects (Bungert et al., 2017; Laakso et al., 2018; Hartwigsen et al., 2015). In this context, we recently established a novel method to localize the origin of the motor evoked potential (MEP) by combining measurements of hand muscle responses at different coil positions and orientations with simulations of the induced E-field (Weise et al., 2020; Numssen et al., 2021b; Weise et al., 2023). In this so-called “regression approach”, a non-linear (sigmoid-like) correlation between the local E-field and MEP is found at the cortical muscle representation within the primary motor cortex (M1). So far, this powerful modeling framework has exclusively been used in the primary motor cortex. The current study explored whether and how this approach may be transferred to the cognitive domain in the healthy human brain. We chose attention as a prototypical function that is relevant to all higher cognitive processes and plays a central role in everyday behavior (Johnson and Proctor, 2004; Schuwerk et al., 2017).

With respect to the underlying neural correlates of attention, it has been demonstrated that different attentional subprocesses are organized in various large-scale networks in the human brain (Corbetta and Shulman, 2002). In particular, two separate networks were identified for visuo-spatial attention, related to the voluntary deployment of attention (i.e., attentional orientation) and the reorientation to unexpected events (i.e., attentional reorientation), respectively (Vossel et al., 2014). Attentional orientation is associated with the dorsal attention network (DAN), which comprises the superior parietal lobule (SPL)/intraparietal sulcus (IPS) and the frontal eye fields (FEF) (Szczepanski et al., 2013). In contrast, the ventral attention network (VAN), including the right inferior parietal lobule (rIPL) and the ventral frontal cortex (VFC), is typically involved in detecting unattended stimuli and triggering attentional reorientation (Corbetta et al., 2008). Previous neuroimaging studies have identified the rIPL as a key region for attentional reorientation, and damage to this area can cause severe attention deficits (Igelström and Graziano, 2017; Numssen et al., 2021a). In this study, we applied TMS to the rIPL while participants were performing a Posner-like attentional task (Posner, 1980). We reasoned that direct modulation of task performance with TMS should provide insights into the underlying causal relevance of specific brain regions for a given task (Walsh and Cowey, 2000).

In summary, we first localized attentional processes with functional neuroimaging at the individual subject level and then applied TMS during the attention task over the individually identified rIPL region. Provided that stimulation of well-localized neural populations would be responsible for observed behavioral effects, correlating the individual E-field strength with those effects (i.e., modulations of reaction times) should identify the areas that are effectively stimulated.

Based on previous TMS studies in the domain of attention (Rushworth et al., 2001; Chambers et al., 2004), our main hypothesis was that interfering with rIPL activity during task performance should selectively modulate behavior during attentional reorientation without affecting attentional orientation. We focused on reaction time (RT) modulation to quantify the causal effects of the TMS-induced E-fields on attentional processes in a continuous manner. The combination of a behavioral experiment with our previously established localization method should provide new insight into the cortical areas that are relevant for attentional processing. A better understanding of the stimulation effects on cognitive functions will guide more effective stimulation protocols for research and clinical purposes.

As a main result, we only found weak correlations between the cortical stimulation strength and reaction time modulation for the attentional task at the individual subject level. We identified and discussed several factors that potentially impeded larger effect sizes. These limiting factors need to be addressed in future applications of a regression-based TMS localization method to fully leverage its potential for causal brain mapping.

## 2. Material and methods

We performed a functional magnetic resonance imaging (fMRI) and a TMS experiment to localize the neuronal populations that are responsible for the TMS effect (Figure 1). In the fMRI experiment, we calculated individual activation maps while participants performed the Posner-like task. In the subsequent TMS localization experiment, 500 TMS trials were performed, while the coil was placed at varying positions near the identified activation peak. The regression approach was performed to identify effective targets for attentional reorientation. Cortical locations with the highest correlation between the induced E-field and the behavioral consequences are assumed to be the optimal TMS target for the perturbation of attentional function.

**Figure 1.**
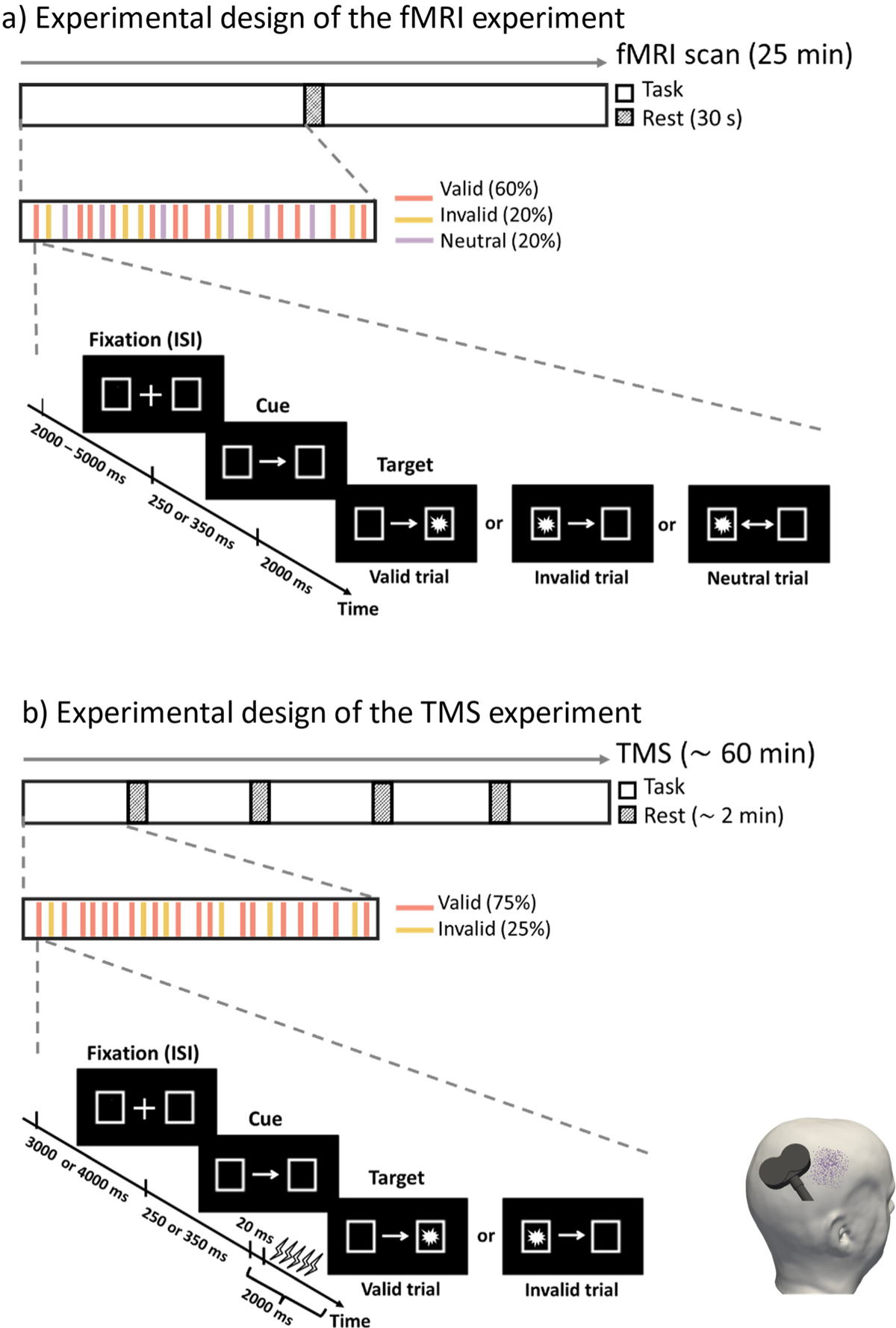
Task paradigm and experimental procedures for the fMRI (a) and TMS experiment (b). A trial consisted of a fixation phase (2-5 s in the fMRI experiment and 3-4 s in the TMS experiment), a cue phase (250 or 350 ms), and a target phase (2 s). Three types of trials (valid, invalid, and neutral) were included in the fMRI experiment, while the neutral condition was excluded in the TMS experiment. A total number of 250 trials were collected in the fMRI experiment. During the TMS experiment, a 5-pulse burst of 10 Hz rTMS was applied 20 ms after the presentation of the target for each trial. 500 TMS trials with random coil positions (represented by violet dots on the head model) and orientations were collected for further analysis.

### 2.1. Participants

Thirty healthy volunteers (15 female, mean age 30.80 ± 5.31 years) were recruited for the fMRI experiment. All participants had normal or corrected to normal vision, and no psychiatric or neurological disorders, or contraindications against TMS or MRI. All participants were right-handed, with a mean laterality index of 90.77 (standard deviation, SD = 10.06) according to the Edinburgh Handedness Inventory (Oldfield, 1971). Written informed consent was obtained from each participant prior to the experiment. The study was approved by the local ethics committee of Leipzig University (ethics number: 371/19-ek). We re-invited seventeen participants (8 female, age 30.12 ± 5.84 years, laterality index 91.24 ± 9.46) to participate in the TMS experiment. Selection criteria were based on fMRI results, with participants being required to show significant activation in the predefined rIPL ROI and a mean error rate in the Posner task below 30%.

### 2.2. Behavioral task

We used an adapted version of Posner’s location-cueing task (Rushworth et al., 2001; Thiel et al., 2004; Numssen et al., 2021a) to trigger orienting and reorienting of spatial attention. The fMRI version of the task contained three trial types: *valid*, *invalid*, and *neutral*, to target different attentional processes (Figure 1). The contrast between invalid and valid trials isolates attentional reorienting processes, and the contrast between valid and neutral trials defines attentional orienting processes/attentional benefits (Grosbras and Paus, 2002; Peelen et al., 2004). During the task, each trial started with the presentation of two rectangular boxes (each size 2.6° of visual angle, with the center situated 8.3° left or right, positioned horizontally) and a fixation cross (size 0.88°) at the center of the screen. To avoid expectancy effects, the duration of fixation (interstimulus interval, ISI) was randomly set to either 2, 3, 4, or 5 s. Subsequently, an arrow was displayed for 250 or 350 ms which served as a cue for the upcoming target location. For valid cues (60 %), the arrow pointed at the side where the target appeared. Participants were trained to orient their attention towards the indicated target position while keeping their fixation at the central fixation cross. For invalid cues (20 %), the arrow pointed to the opposite side of the target, which required participants to reorient their attention to the target side. Neutral cues (20 %) did not contain any information about the target’s location. Finally, a target was presented either at the left or right side (equally distributed) for 2 s. Participants were instructed to identify the positions of the targets by pressing the right/left button using their index and middle fingers as fast and accurately as possible.

We included a total number of 250 trials in the fMRI experiment and 500 in the TMS experiment. The fMRI scan lasted 25 minutes with 150 valid trials, 50 invalid trials, and 50 neutral trials. Participants had a 30 s break in the middle of the task to prevent fatigue (Figure 1a). Before the MRI experiment, participants performed 10 trials as training outside the scanner. In the TMS experiment we specifically focused on attentional reorienting and, thus, only presented valid (75 %) and invalid (25 %) trial types. The ISI was shortened to 3-4 s (Figure 1b). The stimulation lasted approximately 60 min with a short break after every 100 trials. The order of trial types was randomized across participants. Stimuli were administered with Presentation software (v20.1, Neurobehavioral Systems, Berkeley, CA).

### 2.3. MRI data acquisition

Both functional and structural MRI data were collected using a 3 Tesla Siemens Skyra fit scanner with a 32-channel head coil. To segment the main tissues of the head (scalp, skull, gray matter (GM), white matter (WM), cerebrospinal fluid (CSF), and ventricle) for further calculation of the E-field, T1-weighted, and T2* images were acquired with following parameters: T1 MPRAGE sequence with 176 sagittal slices, matrix size = 256 × 240, voxel size = 1 × 1 × 1 mm^3^, flip angle 9°, TR/TE/TI = 2300/2.98/900 ms; T2* with 192 sagittal slices, matrix size = 512 × 512, voxel size = 0.488 × 0.488 × 1 mm^3^, flip angle 120°, TR/TE = 5000/394 ms. The T1-weighted image was also used for neuronavigation during TMS. Diffusion MRI with 88 axial slices, matrix size = 128 × 128, voxel size = 1.719 × 1.719 × 1.7 mm^3^, TR/TE = 80/6000 ms, flip angle 90°, 67 diffusion directions, b-value 1000 s/mm^3^ was acquired for the estimation of the conductivity anisotropy in the WM.

The individual activation map of attentional processing was measured by an event-related fMRI design based on the gradient echo planar (GE-EPI) sequence (60 axial slices, matrix size = 102 × 102, voxel size = 2 × 2 × 2.26 mm^3^, flip angle 80°, TR/TE = 2000/24 ms, 775 volumes). Participants were instructed to perform the Posner task that required spatially congruent button presses in response to visual target stimuli.

### 2.4. MRI data analysis

#### 2.4.1. Preprocessing

MRI data were preprocessed with fMRIPrep 20.1.1 (Esteban et al., 2019), a robust preprocessing pipeline based on Nipype 1.5.0 (Gorgolewski et al., 2011). The individual T1 image was first intensity corrected using N4BiasFieldCorrection (Tustison et al., 2010) and skull-stripped with antsBrainExtraction workflow. Brain tissue segmentation of GM, WM, and CSF was performed on the brain-extracted T1w using FSL FIRST 5.0.9 (Zhang et al., 2001). Brain surfaces were reconstructed using recon-all from FreeSurfer v6.0.1 (Dale et al., 1999), and the brain mask was refined with a custom variation of the Mindboggle method (Klein et al., 2017) to reconcile ANTs-derived and FreeSurfer-derived segmentations of the cortical gray matter. Volume-based spatial normalization to the MNI152Lin standard space was performed through nonlinear registration with the antsRegistration (ANTs 2.2.0).

Functional data were motion-corrected using mcflirt (FSL 5.0.9) (Jenkinson et al., 2002) and slice-time corrected with 3dTshift from AFNI 20160207 (Cox and Hyde, 1997). Distortion correction was performed by 3dQwarp from the AFNI toolbox. The BOLD time series were then co-registered to the T1w reference using bbregister (FreeSurfer), which implements boundary-based registration (Greve and Fischl, 2009), and resampled into the standard MNI152Lin space by antsApplyTransforms (ANTs). Finally, AFNI 3dmerge was used to spatially smooth the functional data with a 6 mm full-width half-maximum Gaussian kernel.

Several motion-induced confounding regressors were also collected based on the preprocessed BOLD: six corresponding rotation and translation parameters, and framewise displacement (FD) (Power et al., 2014).

#### 2.4.2. Activation analysis

Preprocessed data were analyzed with SPM 12 (Wellcome Department of Imaging Neuroscience, London, UK, http://www.fil.ion.ucl.ac.uk/spm). In the first level analysis, a general linear model (GLM) was generated to estimate task-related neural activities. This consists of three regressors of interest: valid, invalid, neutral, and one regressor of no interest: incorrect or missed responses. These regressors were time-locked to the target onset and modeled with a canonical synthetic hemodynamic response function (HRF) with a duration of 0 s. Additionally, six motion regressors and one volume-masking regressor per FD value above 0.9 (Power et al., 2012) were added to regress out motion-induced artifacts. Trials with RT less than 100 ms or greater than 1000 ms were discarded as incorrect responses (Small et al., 2003). After exclusion, the average number of valid trials remaining for analysis was 145 out of 150 (invalid trials: 47 out of 50; neutral trials: 49 out of 50).

On the second level, two t-contrasts were computed to identify areas preferentially engaged in attentional orienting/benefits (valid vs. neutral trials) and reorienting (invalid vs. valid trials), respectively. The contrast of valid vs. neutral trials isolates brain areas activated in trials with targets appearing at the attended position compared to no spatial expectation. The contrast between invalid and valid trials isolates areas activated by targets appearing at the unattended position after a directional spatial expectation was induced by the cue. False discovery rate (FDR) was used to avoid the bias of multiple comparisons (Benjamini and Hochberg, 1995) with *q* < 0.05. Individual peak activation coordinates of invalid > valid on rIPL were extracted to define the cortical area of interest in the subsequent TMS experiment. The Julich brain atlas (Amunts et al., 2020) was used to define the rIPL mask, which comprises one rostral (PGp) and one caudal (PGa) region in the angular gyrus (AG), and five regions of the supramarginal gyrus (SMG) (PFm, PF, PFop, PGcm, PFt).

### 2.5. TMS localization experiment

Seventeen of the thirty fMRI participants underwent an online TMS experiment while performing the Posner-like task (Figure 1b). TMS pulses were applied with coil positions over the larger rIPL area using a MagPro X100 stimulator (MagVenture, firmware version 7.1.1) and an MCF-B65 figure-of-eight coil. Coil positioning was guided by a neuronavigation system (TMS Navigator, Localite, Germany, Sankt Augustin; camera: Polaris Spectra, NDI, Canada, Waterloo). The electromyographic (EMG) signal was amplified with a patient amplifier system (D-360, DigitimerLtd., UK, Welwyn Garden City; bandpass filtered from 10 Hz to 2 kHz) and recorded with an acquisition interface (Power1401 MK-II, CED Ltd., UK, Cambridge, 4 kHz sampling rate) and Signal (CED Ltd., version 4.11).

To determine the optimal stimulation intensity, we manually measured the resting motor threshold (rMT) of the participants’ right index fingers with one surface electrode positioned over the first dorsal interosseous (FDI) muscle belly and one at the proximal interphalangeal joint (PIP). During the rMT measurement, we first positioned the TMS coil over the left-hand knob, which was identified based on established anatomical landmarks (e.g., Diekhoff et al., 2011). The coil was initially placed at 45° to the midline, and then moved around until the optimal coil location and orientation were identified based on the MEP response. The rMT was defined as the minimum stimulator intensity to induce an MEP larger than 50 *μ*V in at least 5 of 10 consecutive trials (Beynel et al., 2019).

500 bursts of rTMS with 5 pulses at 10 Hz each were applied at 100% rMT (Rushworth et al., 2001) during the 500 Posner-like task trials, comprising 375 *valid* trials (75%) and 125 *invalid* trials (25%). The bursts were initiated 20 ms after the target presentation to disrupt attentional processing transiently (Figure 1b). After every 100 trials, a short break of ∼2 min was advised to avoid fatigue. We restricted the area over which the coil centers were located to a circular zone of 3 cm radius around the individual fMRI-derived activation peak (invalid > valid trials) in the rIPL. Stimulation area centers were moved 2 cm anterior/superior in 4 subjects because their activation peaks were close to the occipital or temporal lobe. Coil positions and orientations were randomly selected within the defined circular zone for each burst, to increase electric field variability by minimizing cross-correlations between induced electric fields (Numssen et al., 2021b). The range of coil orientations was limited to 60° (± 30° from the traditional 45° orientation) due to hardware constraints such as the spatial restriction of the navigation system and the obstruction of the coil handle and cable. Note that eleven participants were sampled with a “quadrant mode”: we randomly sampled 100 stimulations for each quadrant of the stimulation area, resulting in 400 trials. The final 100 trials were arbitrarily attributed across the whole stimulation area. To diminish possible sequential effects, we used a “random mode” for the remaining six participants: all 500 trials were randomly sampled throughout the experimental session. Coil positions and corresponding behavioral responses, i.e., reaction time and accuracy, were recorded for each trial.

### 2.6. TMS mapping

#### 2.6.1. Numerical simulations of the induced electric field

Individual head models were reconstructed using the headreco pipeline (Nielsen et al., 2018) utilizing SPM12 and CAT12 (http://www.neuro.uni-jena.de/cat/) for all seventeen subjects. The final head models were composed of approximately 3.4 × 10^6^ nodes and 20 × 10^6^ tetrahedra (average volume: approximately 0.15 mm^3^ in the cortex). Seven tissue types were included with standard conductivity estimates: white matter (*σ_WM_* = 0.126 *S/m*), gray matter (*σ_GM_* = 0.275 *S/m*), cerebrospinal fluid (*σ_CSF_* = 1.654 *S/m*), bone (*σ_B_* = 0.01 *S/m*), skin (*σ_S_* = 0. 465 *S/m*), ventricle (*σ_V_* = 1.654 *S/m*) and eyeballs (*σ_EB_* = 0.5 *S/m*) (Thielscher et al., 2015; Wagner et al., 2004). WM and GM were assigned anisotropic conductivities while the five other tissues were treated as isotropic. Electric fields induced by coil positions of all TMS trials were then computed considering individual head geometry and employing the FEM-based solver implemented in SimNIBS v3.2.6 (Saturnino et al., 2019; Thielscher et al., 2015). See Saturnino et al. (2019) for more details on the numerical simulation.

A refined region of interest (ROI) was defined around the rIPL area based on the Julich brain atlas (Amunts et al., 2020) to improve the numerical resolution around the selected ROI. All analyses were performed on the mid-layer between GM and WM surfaces to avoid boundary effects of the E-field due to conductivity discontinuities.

#### 2.6.2. Regression analysis

The regression approach used in this study is based on a recently developed modeling framework in the motor cortex (Weise et al., 2020; Numssen et al., 2021b; Weise et al., 2023). The principal idea of the regression approach is to combine the outcome of multiple stimulation experiments with the induced E-fields of different coil positions and orientations, assuming that, at the effective site, the relationship between E-field and behavioral performance is stable, i.e., the same electric field strength always evokes the same behavioral output. In particular, the method leverages more information by simultaneously varying both coil positions and orientations exploiting electric field variability to avoid bias towards locations with high E-field magnitudes, such as the gyral crown.

We first discarded TMS trials with RT *<* 100 ms or RT *>* 1000 ms, and those with a coil distance ≥ 5 mm from the skin surface. After applying these exclusion criteria, the average number of trials remaining for analysis was 115 (out of 125) for the invalid condition, and 365 (out of 375) for the valid condition. Subsequently, the valid trials were randomly downsampled to match the number of invalid trials to guard against potential sample-size-dependent effects. In addition, any linear trends of RT over time were removed to mitigate potential learning or fatigue effects.

After data cleaning, we performed Linear regression analysis to identify the neural populations that are causally involved in attentional processing for every element within the ROI (Weise et al., 2020). The linear relationship between E-field magnitude *x_i,j_* of TMS trial *i* (*1* ≤ *i* ≤ *N_TMS_*) at the cortical element *j* (*1* ≤ *j* ≤ *N_elms_*) and the estimated RT *ŷ_i,j_* is calculated:

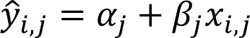

Here, *α* is the intercept and *β* is the slope. We set the constraint of *α* and *β* to (−3000, 3000) and (−1000, 1000), respectively.

The site of effective stimulation can be quantified by the goodness-of-fit (GOF), which would be highest at the cortical site that houses the relevant neuronal populations (Numssen et al., 2021b). We assessed the element-wise GOF by the coefficient of determination *R^2^*:

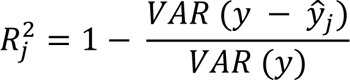

where *y* is the measured RT, and *ŷ_j_* is the estimated RT. The *R^2^* value measures how well the regression model explains the observed data, with higher *R^2^* denoting better fitting results. Note that *R^2^* values can range from negative infinity to 1, where negative values indicate that the chosen model does not follow the trend of the data. Negative *R^2^* values occur when the variance of the residual is greater than the variance of the data, i.e., 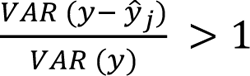.

The regression analysis was applied separately to invalid and valid trials and yielded one *R^2^* score each for valid and invalid conditions per ROI element. We replaced all negative *R^2^* values with 0, as these denote areas of poor fitting and are very small.

To assess the significance of the observed *R^2^* values and determine if they could be explained by chance alone, we carried out a permutation test for each element. This was combined with the FDR analysis to correct for multiple comparisons across different elements. Our permutation test involved randomly shuffling the RT values 1000 times. For each permutation, we calculated the linear regression between the magnitudes of E-field and the shuffled RT values. This process was repeated 1000 times to obtain a distribution of *R^2^* values from chance. We then compared the real *R^2^* value with the 1000 shuffled *R^2^* values. Only real *R^2^* values that exceeded the 95th percentile of the shuffled data were deemed statistically significant. To control the probability of false positives within the ROI, the FDR was employed to keep the false positives rate below 5% for all permutation tests (*q* < 0.05).

We then multiplied the raw *R^2^*value with the slope’s sign to include not only the goodness of fit (*R^2^*) but also the direction of the TMS effect in one metric:

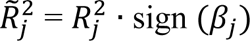

This allowed us to visually differentiate cortical areas of inhibitory TMS effects (positive slope areas) from areas of facilitatory TMS effects (negative slope areas). We assumed that high-frequency rTMS on rIPL will selectively impair performance on invalid trials, resulting in higher *R^-2^* values compared to valid trials.

To generate the group *R^-2^* map, all individual *R^-2^* maps were first mapped to the group averaged brain template, then voxel-wise *R^-2^* values were averaged across subjects.

## 3. Results

### 3.1. Behavioral performance

Behavioral results from the fMRI experiment showed a reliable difference of RT between conditions *(F_2,87_* = 5.38, *p* < 0.01; invalid_fMRI_: 403 *±* 69 ms; valid_fMRI_: 345 *±* 70 ms; neutral_fMRI_: 384 *±* 72 ms). Post-hoc t-tests confirmed a reorienting effect, that is, slower RTs for invalid compared to valid targets (*t_29_* = 15.25, *p* < 0.01). There was no significant difference between neutral and valid or invalid trials (Figure 2a). No significant differences were found in accuracy (*F_2,87_* = 0.92, *p* = 0.40; invalid_fMRI_: 0.95 *±* 0.08, valid_fMRI_: 0.97 *±* 0.06, neutral_fMRI_: 0.95 *±* 0.06).

**Figure 2.**
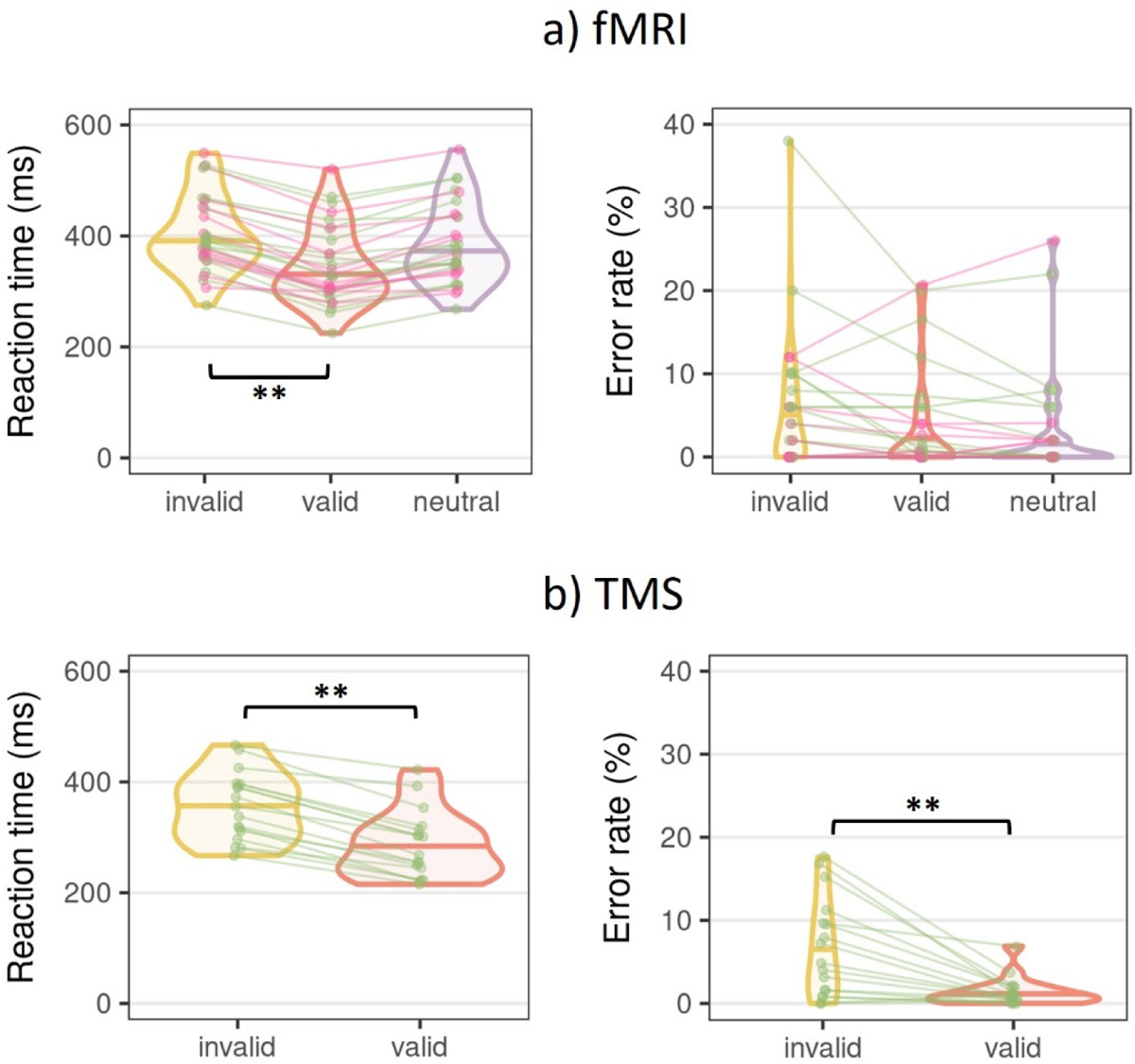
Behavioral results of the Posner task during fMRI and TMS. Reorienting attention (*invalid* trials) was significantly slower than attentional orientation (*valid* trials), both in the fMRI (a) and in the TMS (b) experiment. For task accuracy, significant differences were only present during TMS. ** *p* < 0.01. Green points/lines: subjects who participated in both fMRI and TMS sessions; pink points/lines: subjects who only participated in the fMRI experiment.

In the TMS experiment, we again found a significantly slower response speed for invalid relative to valid trials (*t_16_* = 12.76, *p* < 0.01; invalid_TMS_: 357 *±* 62 ms; valid_TMS_: 286 *±* 61 ms). In line with our hypothesis, task accuracy was also significantly decreased in the invalid condition (*t_16_* = 4.16, *p* < 0.01; invalid_TMS_: 0.93 *±* 0.06; valid_TMS_: 0.99 *±* 0.02) (Figure 2b).

Note that subjects had generally faster RTs in the TMS experiment compared to the fMRI session. These behavioral differences between sessions inside and outside of the MRI scanner might be driven by unspecific distracting factors inside the scanner, such as the supine position and the noise, which could slow down motor execution times and decrease attentional focus inside the MRI scanner (Jamadar et al., 2010; van Maanen et al., 2016; Koch et al., 2003).

### 3.2. fMRI activation

The fMRI results revealed distributed bilateral brain regions from different networks for the reorienting of attention (Figure 3a). These include the ventral attention network (IPL, MFG), the dorsal attention network (SPL, FEF), and the salience network (Insula; anterior cingulate cortex, ACC). In contrast, the default mode network (DMN) (posterior cingulate cortex, PCC; medial prefrontal cortex, mPFC) was deactivated for invalid relative to valid trials. These findings go beyond the classic view of a ventral, right-lateralized reorientation system. Relative to the neutral condition, attentional orienting showed deactivation of both IPS and SPL, but increased activation of mPFC (Figure 3b). All activations are reported at the level of *q* < 0.05 after FDR correction.

**Figure 3.**
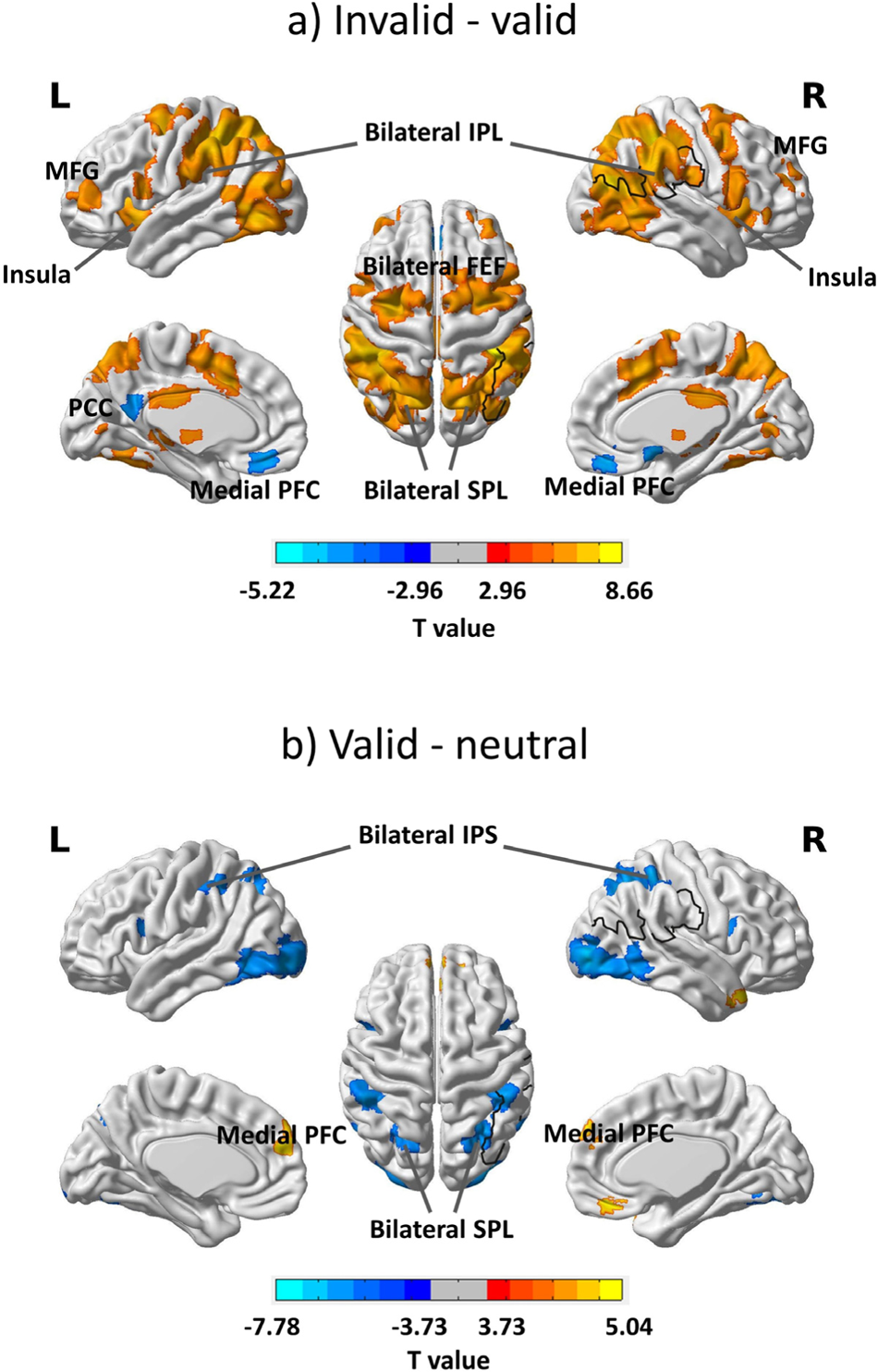
Spatial attention recruits distributed networks: fMRI activation results during attentional orientation and reorientation. (a) Relative to valid trials, the invalid condition exhibits significant activation in both the dorsal and ventral attention networks, as well as the salience network. (b) Attentional orientation resulted in increased activation of the medial prefrontal cortex and deactivation of parietal areas (FDR correction, *q* < 0.05).

### 3.3. TMS mapping results

To elucidate the structure-function relationships in spatial attention, we correlated the magnitudes of the E-field with behavioral performance in the Posner task. Figure 4 displays correlation maps (i.e., *R^-2^*maps) obtained from the TMS mapping experiment, alongside corresponding fMRI activation maps, for all seventeen participants. Warm colors in the *R^-2^*maps indicate positive correlations between E-field and reaction time, while cold colors signify negative correlations. The green background indicates the group-averaged results, while the pink and cyan backgrounds represent the “quadrant sampling mode” and “random sampling mode”, respectively.

**Figure 4.**
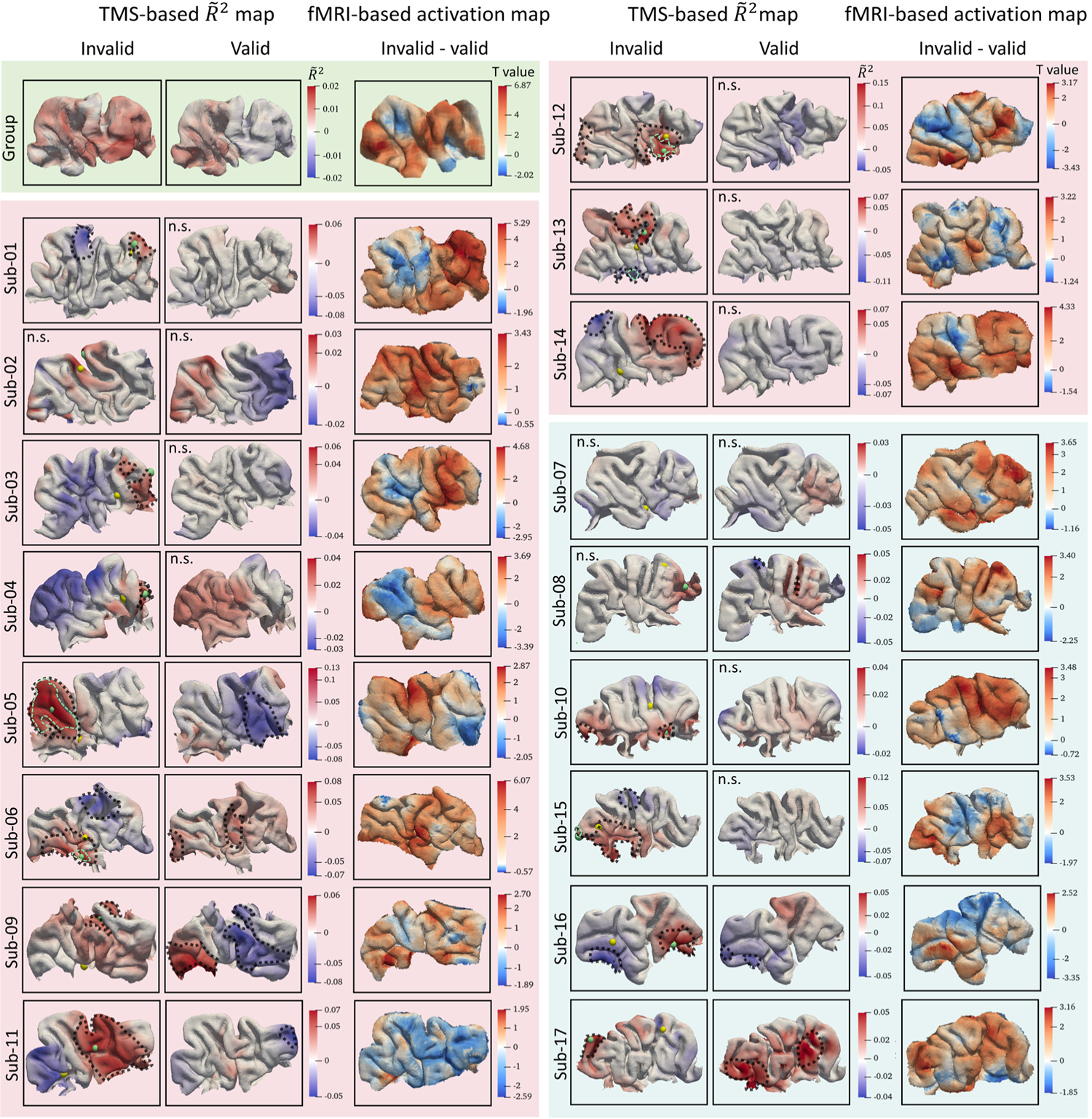
TMS effects on attentional reorientation are subject-specific. This figure shows the group (green background) and individual *R^-2^* maps (pink and cyan background) on the rIPL, as well as the corresponding activation maps. *R^-2^* maps were generated by *R^2^* scores multiplied with the slope’s sign to represent the direction information of the linear correlation. Pink background color codes subjects sampled with “quadrant mode”, while cyan color denotes “random sampling mode” (see text). Yellow spheres in the invalid column represent the fMRI activation peak location of the invalid-valid contrast, whereas green spheres denote the peak *R^-2^*. The black contours marked the brain areas that showed significant *R^2^* values after the permutation test, whereas the light green contours delineate the surviving areas after FDR correction (*q* < 0.05). Note that the *R^-2^* scores were mapped onto the mid-layer surface between white and grey matter whereas the fMRI activation results are mapped onto the cortical grey matter surface. N.s.: not significant after permutation test.

In Figure 5, we present results of a representative subject with moderate fitting results, showcasing the linear regression maps from prominent locations. As shown at three elements (one with the highest GOF of negative linear trend, one on the activation peak, and one with the highest GOF of positive linear trend), the correlation between measured RTs and the computed E-fields did not show a clear S-shaped curve, as MEP data did (Weise et al., 2020; Numssen et al., 2021b).

**Figure 5.**
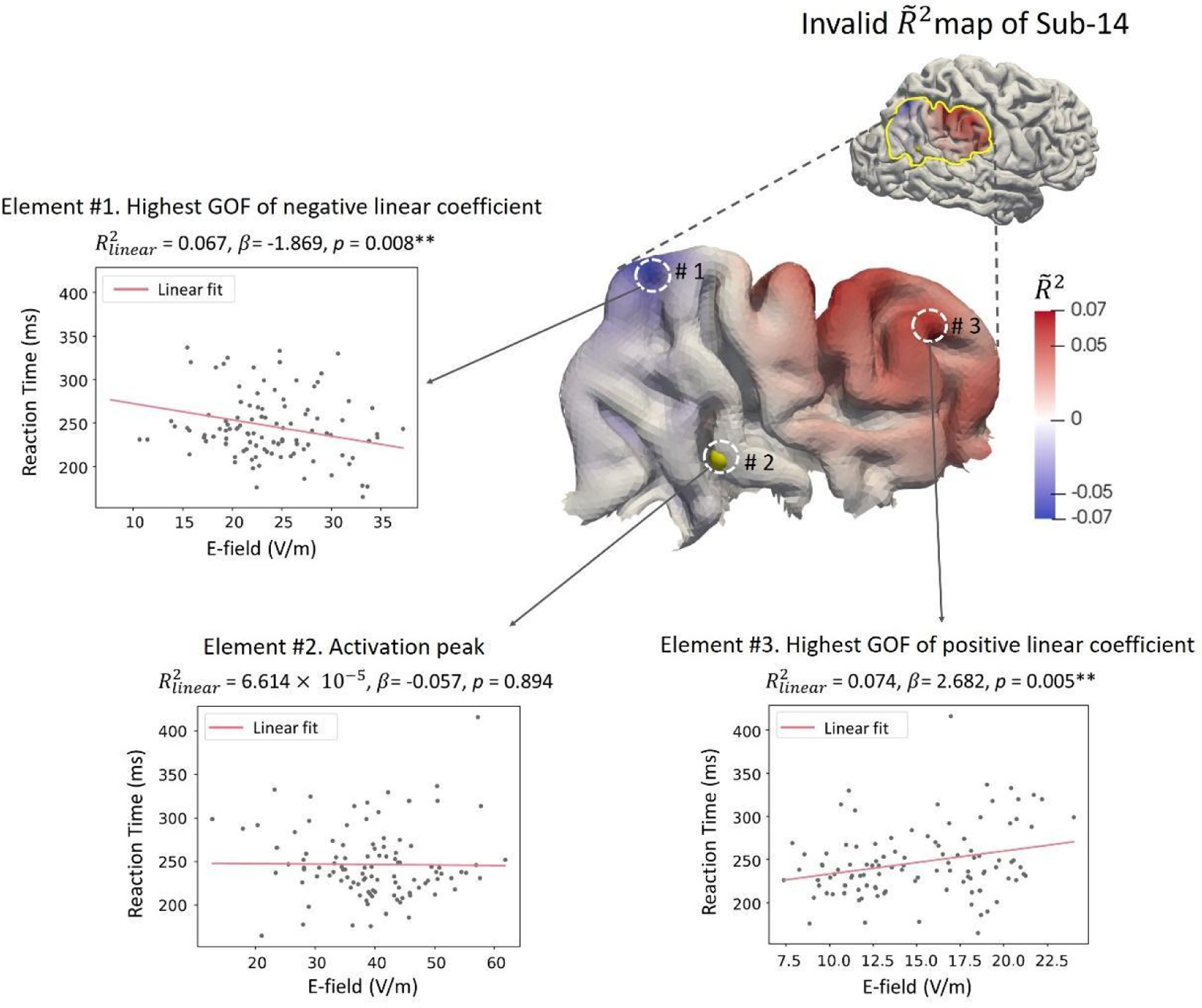
TMS-induced modulation of attentional reorientation from one representative subject with moderate GOF (sub-14). The *R^-2^* map is derived from voxel-wise linear regression of invalid trials, multiplied with the sign of slope to include information about inhibitory (positive slope values) versus facilitatory TMS effects (negative TMS values). The linear fitting lines are shown for 3 elements: #1 with the highest GOF of negative linear correlation, #2 is the fMRI activation peak of reorientation, and #3 with the highest GOF of positive linear correlation. The yellow sphere embedded in the gyral crown marks the location of the fMRI activation peak. *β* denotes the slope of linear regression, and *p* values are the significant level of *β*. **: *p* < 0.01. Grey points: single TMS trials.

At the subject level, most participants showed significant correlations between E-fields and reaction time within the rIPL ROI after permutation test (*p* < 0.05), especially in the invalid condition (Figure 4). The significant brain regions are marked by black contours in Figure 4. After FDR correction, only five subjects (sub-05, sub-06, sub-12, sub-13, and sub-15) retain their significance for the invalid condition. Surviving brain regions are highlighted by light green contours. No significant results were found in the valid condition, implying that TMS selectively modulated attentional reorientation. Notably, the locations of these surviving regions are subject-specific. For instance, in sub-05, the significant region is situated in the angular gyrus, whereas sub-12 shows significant results in the supramarginal gyrus.

Furthermore, to better visualize the pattern differences between the TMS-based *R^-2^* map and the fMRI activation map, we highlighted the locations of both maximum *R^-2^* values (marked by green spheres) and the fMRI activation peak (marked by yellow spheres) across all subjects (Figure 4). The distinct peak locations and patterns between these two maps underscore differences in BOLD-measured neuronal activity and TMS-evoked neuronal effects.

At the group level, the averaged *R^-2^* maps (Figure 4) exhibit values close to zero for both valid and invalid conditions, indicating the absence of a clear-out group-level pattern in the TMS modulatory effect on spatial attention. We then extracted the individual maximum *R^-2^* scores both under the invalid and valid conditions for further comparison. Table 1 shows the maximum *R^-2^* scores for each condition. Given the dominance of positive *R^-2^* values within the group-averaged *R^-2^* map, we only present *R^-2^* scores that show a positive correlation between E-field and RT in Table 1. A Wilcoxon signed rank test revealed a significant difference of the maximum *R^-2^* between the two conditions 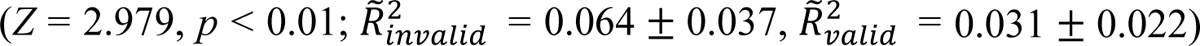 (Figure 6). This indicates that, when compared to the valid condition, the maximum *R^-2^* values are significantly higher in the invalid condition, identifying a stronger disruptive effect of the E-field on the response speed during attentional reorienting.

**Table 1.** Individual activation peaks and TMS results.

| Subject ID | Activation peak | MT | Invalid $\tilde{R}_{max}^2$ | Valid $\tilde{R}_{max}^2$ |
| --- | --- | --- | --- | --- |
| 01 | (56, -37, 34) | 43% | 0.055* | 0.021 |
| 02 | (42, -55, 50) | 39% | 0.031 | 0.024 |
| 03 | (60, -41, 31) | 70% | 0.059* | 0.007 |
| 04 | (54, -33, 34) | 47% | 0.041* | 0.024 |
| 05 | (54, -61, 18) | 57% | 0.127* | 0.079* |
| 06 | (62, -57, 34) | 64% | 0.084* | 0.054* |
| 07 | (52, -53, 16) | 46% | 0.013 | 0.013 |
| 08 | (42, -73, 40) | 43% | 0.030 | 0.048* |
| 09 | (56, -63, 25) | 45% | 0.039* | 0.066* |
| 10 | (58, -47, 36) | 45% | 0.037* | 0.02 |
| 11 | (46, -69, 20) | 42% | 0.073* | 0.036* |
| 12 | (69, -39, 34) | 53% | 0.152* | 0.012 |
| 13 | (52, -51, 31) | 56% | 0.071* | 0.015 |
| 14 | (54, -67, 27) | 38% | 0.074* | 0.008 |
| 15 | (42, -77, 29) | 40% | 0.109* | 0.006 |
| 16 | (56, -67, 34) | 52% | 0.047* | 0.029* |
| 17 | (58, -47, 50) | 44% | 0.045* | 0.058* |
| Group average | (51, -55, 32) | 48% | 0.064 | 0.031 |

Notes: The Activation peak column provides individual fMRI peak coordinates in the rIPL for the invalid-valid contrast in standard MNI space. The individual resting motor threshold (MT) is provided in the percentage of maximum stimulator output (% MSO), and the maximum $\tilde{R}^2$ , both for invalid and valid conditions with positive correlations between E-field and RT. \*: $\tilde{R}_{max}^2$ remains significant after permutation test ( $p < 0.05$ ).

**Figure 6.**
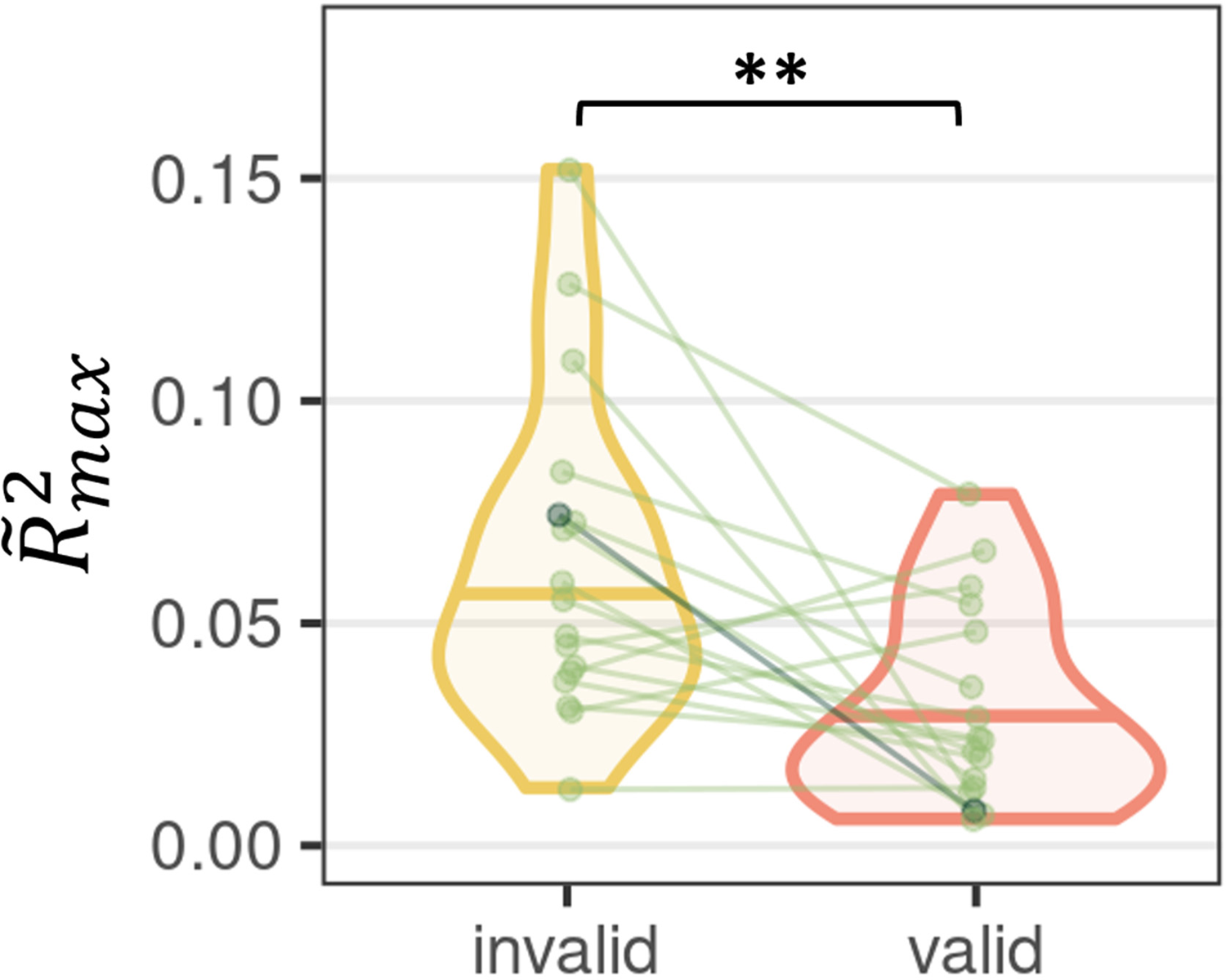
TMS selectively affects attentional reorienting. Invalid trials (reflecting attentional reorienting) were significantly stronger related to the induced E-field than valid trials (reflecting attentional orientation). **: *p* < 0.01. Green points/lines: single subjects. The dark green points/line indicate subject-14, whose detailed *R^-2^* results are shown in Figure 6.

## 4. Discussion

We applied our previously established regression approach to localize the neuronal underpinnings of attentional processes with TMS. In this statistical approach, cortical stimulation exposure across multiple stimulation sites is related to the TMS-induced change in behavior. Here, we studied the relation between the strength of the TMS-induced E-field in the cortex and the inhibitory effects of TMS on spatial attention, to identify relevant cortical regions for attentional processing within the rIPL. We found a weak correlation between E-field strength and attentional performance, highlighting that the translation of the regression approach from motor function to cognitive domains is challenging. At the group level, we observed that the task-specific inhibitory TMS effect was stronger for attentional reorienting relative to attentional orienting (Figure 6). At the single-subject level, the regression analyses identified large individual variability, both for the cortical organization as well as for TMS responsiveness. In addition, we identified differences in BOLD-measured neuronal activity and TMS-evoked neuronal effects. This incongruence highlights the principal distinction between neural activity being correlated with (or maybe even caused by) particular paradigms, and activity of neural populations exercising a causal influence on the behavioral outcome.

### 4.1. fMRI results

The ability to orient attention is a fundamental component of most perceptual-motor processes in everyday life (Natale et al., 2009). Consequently, we took our first step in generalizing the regression method to spatial attention. Spatial orienting can be driven either endogenously, by predictive contingencies, i.e., top-down cues, or exogenously, by unexpected bottom-up signals stemming for example from visual inputs. These processes are accomplished by the interaction of several cortical regions, forming functional networks (Chica et al., 2011). Our fMRI results show that large-scale brain networks were activated during attentional reorienting, including both dorsal and ventral attention networks, which reflects the neural bases of the interplay between these two attention mechanisms (Shulman et al., 2009; Proskovec et al., 2018). One of these regions, the rIPL, is consistently identified as a major network hub in diverse cognitive functions, from bottom-up perception to higher cognitive capacities that are unique for humans (Cabeza et al., 2012; Igelström and Graziano, 2017).

The rIPL is a large region that comprises two major gyri: SMG and AG, separated by the intermediate sulcus of Jensen (Segal and Petrides, 2012). Previous studies suggested that both SMG and AG are critical for attention shifts between visual stimuli (Chambers et al., 2004). A behavioral and functional connectivity-based meta-analysis study further defined the functional topography of rIPL (Wang et al., 2016). This study revealed that the SMG primarily participates in attention, execution, and action inhibition, while the AG is involved in social and spatial cognition. In line with our group activation results, both SMG and AG exhibited significant activation during attentional reorientation, supporting their involvement in basic attention and spatial cognition.

It is worth noting that we found bilateral IPL and VFC activated when participants attempted to process the target at an unexpected location, which challenges the traditional view that the VAN is lateralized to the right hemisphere (Lunven and Bartolomeo, 2017). The comparison of attentional orientation and neutral trials (valid - neutral) allowed us to separate attentional benefits from visual, motor, or other basic cognitive tasks (i.e., valid - baseline), and to explore the influence of cue predictiveness on spatial attention. Previous studies could not provide conclusive explanations for this effect of top-down predictions. Some researchers observed significant activation for valid minus neutral trials in either DAN or VAN (Peelen et al., 2004; Natale et al., 2009), while others found a significant deactivation of the rIPL when participants are orienting their attention to the predictive cues (Doricchi et al., 2010). Our results demonstrate that, when attention was focused on the valid side, bilateral IPS and SPL were suppressed, potentially to prevent reorienting to distracting events. Specifically, activation of medial PFC as a part of DMN may indicate an endogenous focus. These findings provide new insight into the functional contribution of the brain regions involved in spatial attention.

### 4.2. TMS localization

We performed state-of-the-art FEM-based E-field simulations to allow for realistic quantifications of the effect of cortical stimulation on attentional processes. The analysis revealed significant correlations between E-fields and behavioral outcomes in the majority of subjects after the permutation test. However, only five out of seventeen subjects maintained their significance under the invalid condition after FDR correction. This implies that the modulatory effect of TMS on attentional reorientation is weak, exhibiting significant variability across subjects.

Supporting evidence for the specificity of the TMS effect is shown in the significantly stronger relationships observed between E-field exposure and behavioral modulation during attentional reorientation compared to attentional orientation (Figure 6). However, in general, the TMS-induced effect, as measured by the coefficient of determination *R^-2^*, was much lower for attention as compared to the motor domain where the TMS regression approach was previously established (Weise et al., 2020; Numssen et al., 2021b). Lower *R^-2^* values likely reflect the more restricted impact of the E-field on reaction time compared to its effect on muscle recruitment when stimulating the primary motor cortex.

In fact, TMS studies with cognitive paradigms often show high inter-individual variability and results are not always conclusive (Bergmann and Hartwigsen, 2021). Moreover, some cognitive paradigms suffer from relatively low test-retest reliability which may contribute to the strong variability of behavioral results (Hedge et al., 2018). The key problem is that the underlying neural basis of cognitive functions is much more complex and variable than that of eliciting hand muscle twitches (Fetsch, 2016). In our study, in spite of this high variability, we were still able to distill out the TMS effect on attentional reorienting. However, the large amount of residual variance indicates that taking into account additional variables and more complex models (e.g., multivariate regression) is likely to improve accuracy and reliability.

### 4.3. TMS vs. fMRI localization

We paralleled the fMRI-based activation maps with TMS-based cortical mapping to explore the gains of regression-based functional TMS mappings in the domain of spatial attention. The comparison between BOLD-measured neuronal activity obtained from fMRI and neuronal stimulation effects evoked by TMS revealed notable discrepancies. This incongruence highlights the principal distinction between neural activity being correlated with or potentially caused by particular paradigms and the activity of neural populations that exert a direct causal influence the behavioral outcomes. While fMRI provides information about overall neural responses associated with spatial attention, TMS offers insights into the specific and causal effects of targeted neural modulation.

### 4.4. Transferring the TMS regression approach to the study of cognition

The TMS regression approach integrates information from multiple stimulation patterns to map structure-function relationships based on the causal effect of the induced electric field. The main metric to separate cortical locations is the explained variance of the behavioral modulation (*R^-2^*). However, when applied to attentional processes, the explained variance is relatively low. In comparison to the motor domain, where up to 80% of the variance can be explained at the single subject level (Numssen et al., 2021b), only 10% of the individual variance could be explained in the attention domain in the present study. These comparatively low *R^-2^* values for attentional processes probably stem from various sources.

As a first potential explanation for the observed discrepancy between the motor and attention domains, behavioral responses to cognitive tasks intrinsically exert higher variance due to complex processing demands (Hedge et al., 2018) that give rise to confounding factors, such as fatigue or differences in the mental state. Additionally, the organization of the rIPL is challenging to study due to the complex anatomy and highly divergent functional segregation of this cortical region (Williams et al., 2022; Krall et al., 2016).

Another potential factor that may affect the explained variance is the spread of attentional task processing across the cortex. In contrast to the focal representations within the motor system, a multitude of cortical locations may interact with the stimulation effect during attentional processing. The TMS trials with different coil positions/orientations may differentially target cortical sites that have various functions. Some of the varying targets can accumulate and thus exert inhibitory or excitatory effects, others might neutralize the TMS effect. Previous studies confirmed that compared to single-node TMS, concurrent frontal-parietal network TMS showed a reduction of the reorienting effect in the right hemifield (Gallotto et al., 2022). Therefore, network effects should be considered in future studies. One possibility would be the multivariate regression schemes.

Several limitations may have contributed to the limited explained variance. Firstly, we did not adjust the stimulation strength to account for the differences of the cortical depths between the primary motor cortex, where the rMT was assessed, and IPL, where the regression localization approach was performed. Instead, we used the same intensity of 100% rMT as in previous studies (Rushworth et al., 2001) to modulate spatial attention. Ignoring potential individual differences between brain regions may have contributed to the observed variability of the current study. To elaborate on this potential shortcoming, we further computed E-field ratios between the rIPL and the M1 hotspot for all participants. This revealed that the E-field exposure in the rIPL was consistently lower (maximum ratio = 0.95, minimum ratio = 0.25, median ratio = 0.73 across subjects). Subsequently, we conducted Spearman correlation analyses to explore the relationship between the E-field ratio and the maximum *R^-2^* value across participants to further explore if stronger stimulation yielded better functional localization. However, the results did not reach statistical significance (*ρ_invalid_* = 0.18, *p* = 0.48).

Secondly, the current study employed the 5/10 methods to determine the rMT, which is defined as the minimum stimulus intensity required to elicit a peak-to-peak MEP amplitude greater than 50 *μ*V in 5 out of 10 consecutive trials. This rMT determination method has been reported as relatively less reliable than the 10/20 method, which requires eliciting an MEP amplitude greater than 50 *μ*V in 10 out of 20 consecutive trials (Awiszus, 2012). Therefore, the utilization of the 5/10 method could potentially contribute to additional sources of variance. Another limitation of our study is the absence of a sham TMS condition. Consequently, the left part of the input/output line, representing the linear regression between magnitudes of E-field and behavioral outcomes, is missing. This missing information could provide valuable insights into the baseline excitability of the motor cortex. In addition, while we attempted to maximize the range of coil orientations to minimize potential cross-correlations between electric fields, hardware limitations, such as the visible range of positions in the navigation system and coil handle obstruction, limited the true orientation range to 60°.

Finally, it should be noted that the interaction between internal factors such as the current brain state, fatigue, baseline performance level, and external stimulation parameters such as intensity, frequency, and duration is not well understood (Hartwigsen and Silvanto, 2022). Such interactions may induce strong inter-individual variability in response to TMS in studies of cognition. Nonetheless, transferring the TMS regression approach to cognitive domains is promising and will ultimately help to optimize TMS protocols for a wide range of applications. Future work should explore network effects of different TMS protocols, dosing, and individual differences in response to TMS.

## 5. Conclusion

To conclude, we applied a localization approach based on functional analyses of the TMS-induced E-fields and their behavioral modulation in a cognitive domain. With this causal approach, we calculated TMS effects on attentional reorientation and highlighted, both, interindividual variation in cortical organization and differences between fMRI activity and TMS mapping. The results show that E-field modeling can play a valuable role when exploring structure-function relationships with non-invasive brain stimulation methods. We are confident that our approach of combining E-field modeling and behavioral modulation can be further generalized and applied to other functional domains to increase TMS effectiveness and allow its applications at the individual level. To increase the specificity and sensitivity of the method, we suggest developing multivariate regression approaches that account for the recruitment of distributed networks for different cognitive functions. Moreover, alternative readout variables, such as physiological (e.g., heart rate) and electrophysiological (e.g., TMS-evoked electroencephalogram potentials) measures may further increase the information obtained from TMS mapping in cognitive studies.

## Acknowledgments

The authors would like to acknowledge the support from the German Research Foundation (grant no. HA 2899/31-1, HA 6314/9-1, KN 588/10-1 to JH, GH and TRK; and grant no. WE 59851/2 to KW). GH is supported by the Max Planck Society and the European Research Council (ERC-2021-COG 101043747). YJ would also like to thank the China Scholarship Council for its financial support.

